# Optical O_2_ sensors also respond to redox active molecules commonly secreted by bacteria

**DOI:** 10.1101/2022.08.08.503264

**Authors:** Avi I. Flamholz, Samuel Saccomano, Kevin Cash, Dianne K. Newman

## Abstract

From a metabolic perspective, molecular oxygen (O_2_) is arguably the most significant constituent of Earth’s atmosphere. Nearly every facet of microbial physiology is sensitive to the presence and concentration of O_2_, which is the most favorable terminal electron acceptor used by biological organisms and also a dangerously reactive oxidant. As O_2_ has such sweeping implications for physiology, researchers have developed diverse approaches to measure O_2_ concentrations in natural and laboratory settings. Recent improvements to phosphorescent O_2_ sensors piqued our interest due to the promise of optical measurement of spatiotemporal O_2_ dynamics. However, we found that our preferred bacterial model, *Pseudomonas aeruginosa* PA14, secretes more than one molecule that quenches such sensors, complicating O_2_ measurements in PA14 cultures and biofilms. Assaying supernatants from cultures of 9 bacterial species demonstrated that this phenotype is common: all supernatants quenched a soluble O_2_ probe substantially. Phosphorescent O_2_ probes are often embedded in solid support for protection, but an embedded probe called O_2_NS was quenched by most supernatants as well. Measurements using pure compounds indicated that quenching is due to interactions with redox-active small molecules including phenazines and flavins. Uncharged and weakly-polar molecules like pyocyanin were especially potent quenchers of O_2_NS. These findings underscore that optical O_2_ measurements made in the presence of bacteria should be carefully controlled to ensure that O_2_, and not bacterial secretions, is measured, and motivate the design of custom O_2_ probes for specific organisms to circumvent sensitivity to redox-active metabolites.

**Importance:** When they are closely-packed, as in biofilms, colonies, and soils, microbes can consume O_2_ faster than it diffuses. As such, O_2_ concentrations in natural environments can vary greatly over time and space, even on the micrometer scale. Wetting soil, for example, slows O_2_ diffusion higher in the soil column, which, in concert with microbial respiration, greatly diminishes [O_2_] at depth. Given that variation in [O_2_] has outsized implications for microbial physiology, there is great interest in measuring the dynamics of [O_2_] in microbial cultures and biofilms. We demonstrate that certain classes of bacterial metabolites frustrate optical measurement of [O_2_] with phosphorescent sensors, but also that some species (e.g. *E. coli*) do not produce problematic secretions under the conditions tested. Our work therefore offers a strategy for identifying organisms and culture conditions in which optical quantification of spatiotemporal [O_2_] dynamics with current sensors is feasible.

## Observation

The O_2_/H_2_O redox couple (E’° ≈ +800 mV) is the most favorable terminal electron acceptor used by biology. Indeed, aerobic microbes typically regulate their metabolisms to use O_2_ before any other terminal acceptor (1), even those with very favorable midpoint potentials like nitrate (NO_3_^-^ →NO_2_^-^, E’° ≈ +370 mV). Today, O_2_ is abundantly produced by photosynthetic organisms, leading to an atmospheric partial pressure of ≈21% and equilibrium aqueous concentrations of ≈200-400 μM (2). This enormous reservoir of oxidant powers the complex multicellular organisms and vibrant ecosystems living near Earth’s surface.

Yet a wide variety of natural environments, ranging from soils to the insides of animals, are characterized by very low O_2_ concentrations ([O_2_]). Organisms endemic to such environments are termed “functional anaerobes” as they often rely on enzymes that are irreversibly deactivated by oxygen and cannot grow in high O_2_ settings. This toxicity is due to the reactivity of O_2_ itself as well as reactive oxygen species (e.g. H_2_O_2_ and O_2_^-^) that are produced from O_2_ through biotic and abiotic mechanisms (3). O_2_-impacted physiology also matters practically: low O_2_ correlates with antibiotic tolerance (4).

Microbiologists typically categorize environments as “oxic,” “anoxic,” or “hypoxic,” but natural environments are heterogeneous and often-characterized by intermediate and fluctuating O_2_ levels. Along the murine intestinal tract, for example, [O_2_] can range from sub-ambient in the stomach (≈50 μM) to near anoxia (< 1 μM) in cecum (5). This difference is consequential: microbes express alternative respiratory pathways when [O_2_] is depleted (1). Similarly, intestinal O_2_ levels decrease substantially when respiratory metabolism is stimulated by addition of glucose (6), so the O_2_ level that microbes experience depends on the balance of inflow and local metabolism.

When they are densely-packed in environments like biofilms, colonies, or soils, microbes can consume O_2_ faster than it diffuses, generating sub-oxic microenvironments that affect metabolism and growth (7). Spatial transcriptomic data suggests that such microenvironments induce large changes in gene expression (8). We might intuit that microbes tune their gene expression, metabolism, and physiology in response to the local environment, yet bacterial gene expression is noisy (9) and often imperfectly-optimized for growth and survival even in batch culture (10). We would like to understand how bacteria adapt to communal life in spatially-heterogeneous settings, but our inability to measure the chemical microenvironment limits our capacity to interpret the “omics” data already collected.

Given its centrality to metabolism and physiology, there is great interest in measuring [O_2_] in natural environments (11) and laboratory experiments (12–15). Bulk O_2_ concentrations can be measured via several methods (16). However, few of these approaches can measure spatiotemporal O_2_ dynamics to characterize chemical microenvironments (11, 17, 18). Planar optodes are constructed by coating a surface with a phosphorescent O_2_-sensing dye over which a semi-permeable polymer matrix is applied as insulation. As O_2_ quenches dye phosphorescence, the optode can be calibrated by measuring phosphorescence intensity or lifetime in known O_2_ concentrations (17, 19).

In principle, these dyes could be used in solution to measure [O_2_] at high spatiotemporal resolution. However, O_2_-sensing dyes are also quenched by molecules other than O_2_ (20). This problem has motivated engineering of sensors protected from spurious interactions by chemical modification (20, 21) or polymer encapsulation (5, 13). Impressed by recent measurements of spatiotemporal O_2_ dynamics in rodent brains (21) and intestines (5), we attempted similar measurements in *Pseudomonas aeruginosa* (PA14) biofilms. As PA14 is known for its secretion of many small molecules including redox-active phenazines (22), we first tested whether culture supernatants quench a model soluble O_2_ sensor, RTDP (17, 23).

We grew PA14 in a minimal medium where it reliably produces the phenazine pyocyanin (PYO) and found that RTDP was strongly quenched by filtered supernatants. Undiluted supernatants reduced fluorescence by ≈50% (Figure 1A), similar to ambient levels of O_2_ (24). Quenching of RTDP by organics is well-documented (23) and could be due to several chemical mechanisms (25). We therefore proceeded to test a “protected” sensor, O_2_NS, composed of an O_2_-sensitive platinum-bound porphyrin encapsulated in a spherical ≈200 nm PVC matrix (13). Neat PA14 supernatants quenched O_2_NS fluorescence by ≈40% (Figure S3). We quantified the concentration-dependence of quenching as an effective Stern-Volmer constant 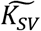 (19); larger values indicate greater quenching (Figure 1C-D).

**Figure 1:**
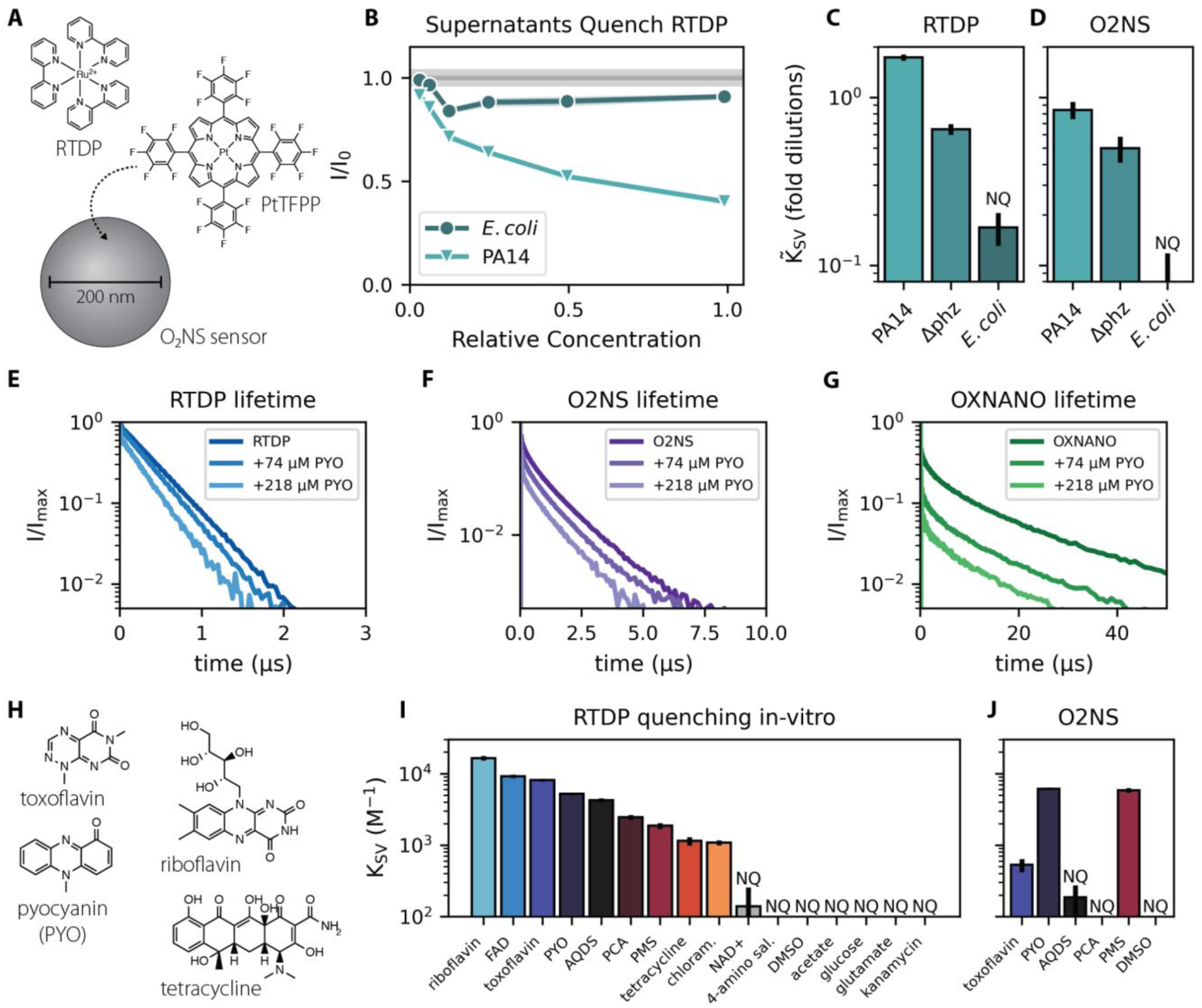
Bacteria secrete soluble quenchers of phosphorescent O_2_ sensors. (A) Tris(2,2’- bipyridyl)dichlororuthenium(II), or RTDP, is a soluble phosphorescent ruthenium complex that can be used as an O_2_ sensor (17). Platinum (II) meso-tetra(pentafluorophenyl)porphine, or PtTFPP, is the O_2_-sensing component of O_2_NS nanosensors, which encapsulate the Pt-porphyrin with a reference dye in a spherical ≈200 nm PVC matrix (13). (B) Strains were grown in a minimal medium where *P. aeruginosa* (PA14) reliably produces pyocyanin (26). Serially-diluted PA14 supernatants quenched RTDP in a concentration-dependent manner. I/I_0_ denotes fluorescence normalized to no quencher, and the width of the horizontal gray bar gives the standard deviation of RTDP intensity in culture media. (C-D) Quantification shows that PA14 supernatants quench RTDP and O_2_NS, and that quenching is partially due to secreted phenazines. The concentration-dependence of quenching was fit to a Stern-Volmer model (19) where 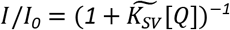. Quencher concentration [Q] has units of (fold dilutions)^-1^ here. 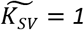 implies I/I_0_ = 0.5 when [Q] = 1, i.e. for an undiluted supernatant; larger 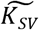 imply stronger quenching. Deletion of phenazine biosynthesis genes (PA14 Δphz) reduced but did not eliminate quenching, and *E. coli* supernatants quenched both sensors far less than PA14. (E-G) Phosphorescence lifetime measurements of RTDP, O_2_NS and a commercial O_2_ sensor, OXNANO, indicate that the phenazine pyocyanin (PYO) is a strong concentration-dependent quencher of all three sensors. Panels give normalized intensities (I/I_max_). (H) A subset of biological molecules tested for RTDP quenching. Panel (I) gives the results of serial dilution measurements akin panel B. Here *K_sv_* has M^-1^ units. Redox active molecules capable of spontaneous electron transfer were stronger quenchers. (J) O_2_NS was quenched by many of the same molecules, but encapsulation appears to protect against quenching by polar and charged species like toxoflavin and PCA. Error bars mark a 95% confidence interval. ‘NQ’ denotes ‘non-quencher’ where the fit to a Stern-Volmer model was poor or quenching was too weak to quantify (Methods, Figure S1). Abbreviations: flavin adenine dinucleotide (FAD), 9,10-Anthraquinone 2,6-disulfonic acid (AQDS), phenazine 1-carboxylic acid (PCA), phenazine methosulphate (PMS), chloramphenicol (chloram.), nicotinamide adenine dinucleotide (NAD+), 4-amino salicylate (4-amino sal.), dimethyl sulfoxide (DMSO).

Photochemical electron transfer is one mechanism by which small molecules quench phosphorescence (25). We hypothesized that quenching is due to secretion of redox-active molecules like phenazines, which is common in *Pseudomonas* species (22). Consistent with this hypothesis, supernatants from a strain lacking phenazine biosynthesis (PA14 Δ*phz*) grew to similar densities as wild-type, but quenched RTDP and O_2_NS to a lesser extent (Figure 1B-C). Nonetheless, PA14 Δ*phz* supernatants still quenched both sensors substantially more than *E. coli* supernatants, with fit PA14 *Δphz* 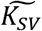 being ≈5-fold larger for RTDP and ≈30-fold larger for O_2_NS. Accordingly, we conclude that PA14 secretes more than one class of molecules that quench phosphorescent O_2_ sensors.

As PA14 makes substantial PYO in our minimal medium (26), we used lifetime spectroscopy to verify that PYO quenches three distinct phosphorescent O_2_ sensors *in vitro:* RTDP, O_2_NS, and a commercial sensor called OXNANO (Figure 1E-G). We next tested a variety of small molecules with diverse biological functions in a serial dilution assay. Redox-active molecules quenched RTDP while controls lacking spontaneous redox activity did not (Figure 1I). As these serial dilution experiments were conducted on the benchtop, we assumed that redox active molecules were predominantly oxidized.

We then assayed O_2_NS with a subset of the molecules considered in Figure 1I. Despite large RTDP quenching constants, molecules carrying net charge at neutral pH (PCA, AQDS) did not quench O_2_NS and were not differentiable from the negative control, DMSO (Figure 1J). This likely reflects the low solubility of charged molecules in the hydrophobic PVC matrix of O_2_NS particles. In contrast, PYO and PMS, which are uncharged and have little polar surface area, quenched O_2_NS and RTDP with large associated *K_sv_* values. Toxoflavin is uncharged but has a relatively larger polar surface area, which may explain its ≈20 fold lower *K_sv_* for O_2_NS.

To understand whether bacteria commonly secrete quenchers of phosphorescent O_2_ sensors, we tested supernatants from 7 additional bacterial species selected based on their ability to grow in our minimal medium supplemented with amino acids. All supernatants quenched RTDP measurably (Figure 2A). O_2_NS was also quenched by several culture supernatants, but less strongly than RTDP (Figure 2B & Figure SX). Despite the change in growth media, *E. coli* supernatants did not quench O_2_NS.

**Figure 2:**
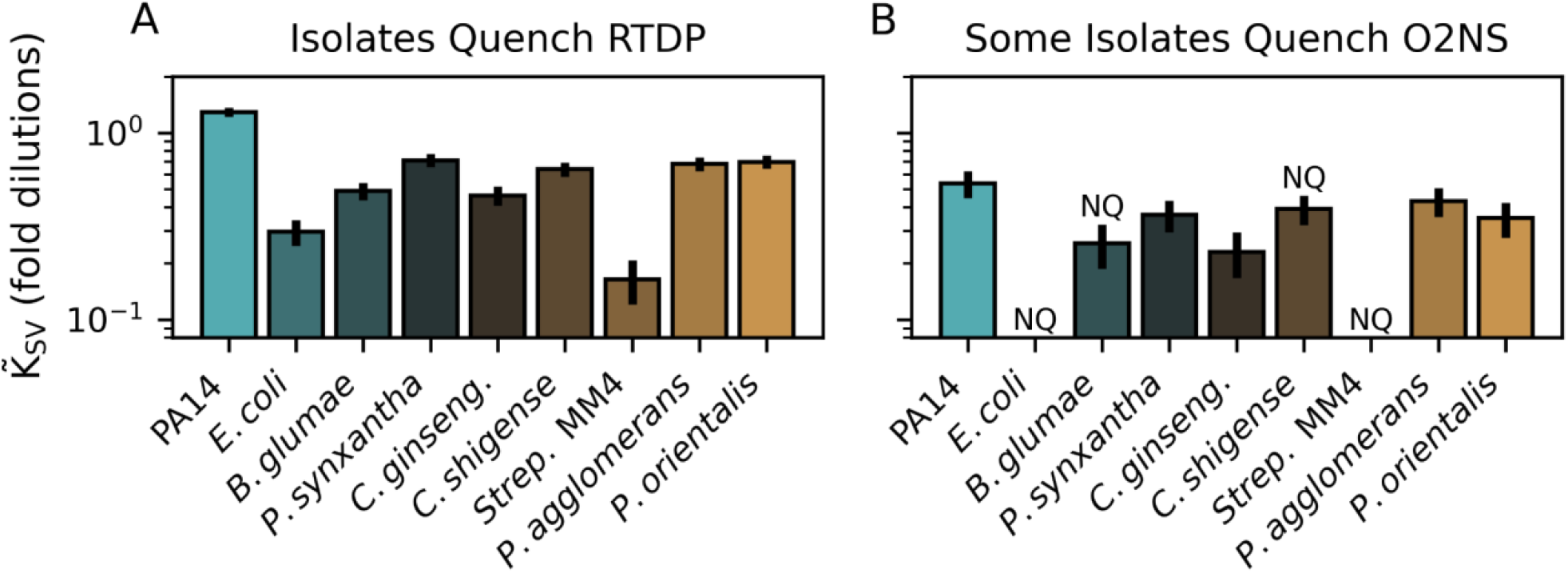
Bacteria commonly secrete soluble quenchers of phosphorescent O_2_ sensors. (A) Supernatants from 7 additional bacterial strains were screened for RTDP quenching activity after growth in a glucose minimal medium supplemented with amino acids. All supernatants quenched RTDP measurably, but to varying degrees. (B) Screened supernatants quenched O_2_NS to a lesser extent. Error bars give 95% confidence intervals. ‘NQ’ denotes ‘non-quencher’ where the fit to a Stern-Volmer model was poor or quenching was too weak to quantify (Methods, Figure S1). Strains are described in Table S1.

## Conclusions

Motivated by the need to measure chemical microenvironments to better understand microbial physiology at the microscale, we attempted to make optical measurements of O_2_ dynamics in the presence of bacterial secretions. We found that model phosphorescent O_2_ sensors were substantially quenched by supernatants from several bacterial species (Figure 1). Quenchers in *P. aeruginosa* PA14 supernatants included, but were not limited to, phenazines (Figure 1B-C). Encapsulating the O_2_-sensitive molecule in a protective matrix, as in O_2_NS, reduced quenching but did not eliminate it in most cases

(Figure 1D and F). *In vitro* experiments indicated that quenching was due to interactions with redox-active secreted molecules like pyocyanin (Figure 2). Indeed, the pyocyanin concentration can surpass 100 μM in PA14 supernatants (22); such concentrations quenched O_2_NS by ≈40%, comparable to ≈30 μM O_2_ (13). Several published works report optical O_2_ quantifications using similar sensors in the presence of growing bacteria (12–15). Such measurements typically rely on calibration curves measured in the absence of cells (12, 13, 15), or in the presence of unknown concentrations of bacterial secretions (14). Our results demonstrate that additional controls are required to demonstrate that the bacteria under study do not produce spurious quenchers in the chosen culture conditions.

With some exceptions (24), RTDP is not usually used to measure [O_2_] in the presence of living cells, likely due to the potential for spurious quenching. Rather, we used RTDP to demonstrate that microbes commonly secrete quenchers (Figure 1D) and intended the RTDP-O_2_NS comparison to illustrate the protective effect of the polymer matrix, which is substantial (Figures 1D-E), but depends on quencher chemistry (Figures 2C & S8). These results indicate that diverse bacteria secrete small-molecule quenchers that are likely redox-active (Figure 1I). We therefore suggest that RTDP might be used to identify copious producers like PA14. Notably, *E. coli* supernatants did not quench O_2_NS (Figure 1C and E), indicating that O_2_NS and similar sensors might succeed in measuring spatiotemporal [O_2_] dynamics in the presence of growing *E. coli* cells. Our protocol for using serial dilutions to estimate supernatant quenching (*Methods* and Figure S1) therefore offers a road-map for identifying mutually-compatible organisms, growth media and O_2_ sensors for optical characterization of the chemical microenvironment. Towards the goal of mapping spatiotemporal variation in O_2_ concentrations in diverse systems, our work spotlights the opportunity and need to design O_2_ sensors appropriate for specific applications.

## Materials and Methods

### Strains and culture conditions

All strains were grown in a glucose minimal medium (GMM) comprising 10 mM glucose 50 mM KH2PO4/K2HPO4 (pH 7.2), 42.8 mM NaCl, 9.35 mM NH4Cl, 1 mM MgSO4 and a trace elements solution. The medium was prepared by autoclaving all the components together, except for the glucose and the 1000× trace elements stock solution, which were sterilized through filtration and added after autoclaving (26). For the experiments reported in Figure 1E-F, 1× MEM Amino Acids solution (MilliporeSigma, Cat. No. M5550) was added so that diverse strains could be grown in the same media. For experiments focused on fast-growing strains (Figure 1A-D) strains were pre-grown to saturation in 5 ml GMM (≈24 hours) and back-diluted by adding 200 μl saturating culture to 5 ml fresh GMM. Experimental cultures were grown for 24 hours before collecting supernatants (see below). For experiments including slower-growing strains (Figure 1E-F) pre-cultures were grown for ≈60 hours, back-diluted and allowed to grow 48 hours before collecting supernatants. Bacterial strains are described in Table S1.

### Chemical stocks

Pyocyanin (PYO) was synthesized from phenazine methosulphate (PMS) as described previously (26) and a 7.5 mM stock was prepared in 20 mM HCl. Unless otherwise noted, other chemical stocks were prepared at 10 mM in a filter-sterilized 25 mM HEPES buffer (pH 7) and stored at 4 °C between experiments. The following chemicals were sourced from Sigma Aldrich: NAD+ (β-nicotinamide adenine dinucleotide hydrate, N1511), FAD (flavin adenine dinucleotide disodium salt hydrate, F6625), riboflavin (R-4500), L-glutamic acid monosodium salt hydrate (G1626-100G), sodium 4-aminosalicylate dihydrate (A-3505) and tetracycline (T-3383). 1 M D-glucose (X) and acetate (sodium acetate trihydrate; Fischer Scientific S209-500) were prepared in milliQ water and diluted to 10 mM in buffer. PMS (Alfa Aesar, H56718) stocks were prepared fresh due to its known photoconversion to PYO. Toxoflavin (MedChemExpress) was dissolved in dimethyl sulfoxide (DMSO, Macron Fine Chemicals) to make a 10 mM stock solution (26). Lab stocks of kanamycin sulfate (Gibco 11815-024, 103 mM in water) and chloramphenicol (Sigma C0378, 77.3 mM in ethanol) were diluted to 10 mM in HEPES buffer. 500x RTDP (100 mg/ml Tris(2,2’-bipyridyl)dichlororuthenium(II) hexahydrate, Sigma Aldrich 544981) stocks were prepared in milliQ water. Multiple O_2_NS batches were prepared on the same date as previously reported (13) and pooled into a single stock that was used for all experiments.

### Measurement of supernatant quenching by serial dilution

Cultures supernatants were collected by centrifugation of 1-2 ml of saturating culture (6 min. at 5000g) in a benchtop centrifuge (BRAND). 500 μl of culture supernatant was filtered through 0.22 μm sterile spin filters (Costar Spin-X, Cat. No. 8160), which were centrifuged for 2 min. at 10000g. Filtered supernatants were then twofold serially diluted 3-5 times in 100 μl volume in a 96 well plate (BRANDplates 781671). Wells containing 100 μl of fresh culture media were included as blanks. Optical measurements were then conducted using a Tecan Spark 10M plate reader. Absorbance spectra were recorded to verify dilution and fluorescence measurements were taken to quantify the baseline (“pre-addition”) fluorescence of each supernatant. RTDP was excited at 450 nm and fluorescence was monitored at 625 nm; for O_2_NS experiments, both dyes were excited at 450 nm, PtFPP was monitored at 650 nm and DiA 585 nm (13). After baseline measurements, the appropriate sensor (RTDP or O_2_NS) was added to all wells and fluorescence was measured again (“post-addition”). For O_2_NS, 11 μl stock was added to each well after which well contents were mixed by pipetting and 11 μl was withdrawn to conserve the total volume. In early experiments (Figure 1A-D), 1 μl of 100x RTDP was added to each well. In later experiments (Figure 1E-F) 11 μl of 10x RTDP was added and withdrawn as described for O_2_NS samples. Culture densities were recorded in 5x dilution at 500 nm (Beckman Coulter DU 800). Strains were grown in biological duplicate and each supernatant was assayed in technical duplicate. The procedure for fitting quenching constants (*K_sv_*) from these data is described below.

### Measurement of pure chemical quenching by serial dilution

Stocks of pure chemicals were diluted to a defined concentration and then twofold serially diluted five times in 100 μl volume in a 96 well plate (BRANDplates 781671). The starting concentrations were as follows: 10 mM acetate and glucose, 500 μM 4-amino salycilate, AQDS, FAD, glutamate, NAD+, PCA, PMS, tetracycline and toxoflavin, 502.5 μM chloramphenicol and PYO, 504.7 μM kanamycin, and 200 μM riboflavin. Wells containing 100 μl of buffer included as blanks. Optical measurements were conducted in a Tecan Spark 10M plate reader as described above, i.e. first recording baseline absorbance spectra and fluorescence levels and then recording fluorescence again after sensor addition (RTDP or O_2_NS). This procedure is diagrammed in Figure S1. Excitation and emission parameters are given above and the procedure for fitting quenching constants (*K_sv_*) is described in the supplement.

### Fitting supernatant quenching from serial dilution experiments

The Stern-Volmer plots in Figures S2, S3, SX, and SX plot sensor I0/I against quencher concentration [Q]. Stern-Volmer predicts a linear relationship of I0/I = 1 + *K_sv_* [Q]. I0 is sensor fluorescence intensity in the absence of any quencher (i.e. in buffer), which was quantified from the post-addition fluorescence of control wells. Since the supernatants and pure chemicals used were often fluorescent, sensor fluorescence in the presence of quencher (I) was calculated from the dilution series by subtracting the rescaled baseline measurement from the post-addition fluorescence. Rescaling was necessary because the pre-addition sample was typically diluted by ≈10% by the addition of the sensor in order to maintain a constant sample volume and path-length. For a 10% dilution, then, I = Ipost - Ipre * 0.9. As 11 ul was added to a 100 ul in most experiments, actual dilution factors were (1-11/111) ≈ 0.9. Given this ≈10% dilution, quencher concentrations were also rescaled before generating Stern-Volmer plots and fitting quenching constants. That is, [Q] = [Qpre]*0.9 for the example of 10% dilution. We then used weighted linear least squares fitting (Python statsmodels package) to fit I0/I to a linear function of [Q]. Data points were weighted by estimated accuracy (1/variance) inferred by error propagation through blanking and ratio calculation. The slope of this fit was taken to be *K_sv_*. Symmetric 95% confidence intervals on *K_sv_* were calculated using the same package. Chemicals and supernatants were marked ‘NQ’ for ‘non-quencher’ if [Q] was poorly correlated with I0/I (Pearson *R* < 0.4) or if the magnitude of quenching overlapped with the variability measured for the unquenched sensor. This analysis pipeline is diagrammed in Figure S1. All source code for data processing and figure generation can be found at github.com/flamholz/secretions_quench_O2sensors.

### Phosphorescence lifetime measurements

All lifetime measurements were conducted in a 25 mM HEPES buffer (pH 7.0). Buffer was air-equilibrated by bubbling house air for 1 hr. RTDP stock was diluted to 40 μM for measurement and O_2_NS was diluted tenfold from stock. A 0.5 mg/ml stock of OXNANO (PyroScience) was prepared and then diluted tenfold into aerated HEPES buffer for measurement. All measurements were conducted in a custom-built lifetime fluorometer in the Caltech Beckman Institute Laser Resource Center. Lifetimes were first measured in 2 ml volumes in quartz cuvettes without PYO to choose measurement parameters and establish a baseline lifetime. All sensors were excited at 355 nm; RTDP and O_2_NS were monitored at 650 nm, OXNANO at 750 nm. PYO stock (7.5 mM) was then added to the cuvette in defined volumes - 2 μl, 18 μl, 20 μl, 20 μl - and lifetimes were measured between additions. Cuvettes were stirred by a magnetic stir-bar during measurement.

## Acknowledgments

Thanks to Chelsey VanDrisse for supplying pyocyanin, Andrew Babbin for OXNANO beads, Lucas Meirelles and John Ciemniecki for assistance with toxoflavin and *B. glumae* cultivation. Thanks to Josh Goldford, Darcy McRose, Georgia Squyers, and Lev Tsypin and for useful conversations. This investigation was aided by a Postdoctoral Fellowship from The Jane Coffin Childs Memorial Fund for Medical Research (to A.I.F.) and NIH grants (1R01AI127850-01A1 and 1R01HL152190-01) to D.K.N as well as the US Department of Energy (DOE) Office of Science, Office of Biological and Environmental Research Bioimaging Science Program under subcontract B643823 (to KJC) and the LLNL 3DQ Microscope Project, SCW1713.

**Supplemental Figures and Tables**

**Table S1: bacterial strains used in this study.**

**Figure S1: diagram of serial dilution method for quantifying quenching.** The approach is diagrammed here for culture supernatants, but an analogous approach was used for pure chemicals. In the latter case quencher concentrations are known and used to fit *K_sv_* in M^-1^ units. (A) Culture supernatants were harvested by centrifugation and filtering after which they were twofold serially diluted in a 96-well plate. (B) Biological duplicate supernatants were serially diluted in technical duplicate and control wells containing only culture media were included in all experiments. (C) A measurement was taken prior to sensor addition to quantify the baseline fluorescence of supernatants as well as culture media for blanking. Sensor was then added in a defined concentration and another measurement was taken. (D) When the supernatants are themselves fluorescent (e.g. blue line) our expectation is that, after sensor addition, a non-quenching supernatant should yield fluorescence intensity equal to the weighted sum of the baseline (supernatant alone) and the sensor alone in buffer/media (gray line). Weighting accounts for ≈10% dilution of supernatant upon sensor addition. The expected sum is given as the dashed purple line. In the presence of quenching, measured fluorescence intensity will fall beneath this expected value, with the exact curve depending on the degree of quenching (*K_sv_*, solid purple lines). (E) To analyze a post-sensor-addition measurement like the curve in purple, we first subtract off baseline (pre-addition) measurement to estimate I and compare it to I_0_ (sensor alone in media/buffer, grey line). Since there is variability in the blank (buffer alone), the pre-addition baseline (blue line), and the reference measurement of the sensor in buffer (grey line), our code propagates the observed uncertainties in these values through the calculation of I_0_/I. The Stern-Volmer curve (inset) is then used to fit *K_sv_* by fitting I_0_/I to a linear function of [Q], here the relative quencher concentration, but absolute concentrations are used in experiments with pure quenchers. In this example we assumed *K_SV_* = 1.0 and 3% normally-distributed measurement error in order to generate the purple curve. Note that the 95% confidence interval on the fit *K_sv_* includes the input value. In cases where I_0_/I was poorly correlated with [Q] (R < 0.4), where the projected quenching of neat supernatant overlapped with the observed variation in signal from unquenched sensor in buffer, or where the projected effect of 1 M pure quencher overlapped with the observed variation in unquenches sensor fluorescence.

**Figure S2: RTDP Stern-Volmer plots for the strains presented in Figure 1C.** In addition to the strains presented in the main text, we tested supernatants from two PA14 mutants - a double siderophore knockout PA14 Δ*pvdA* Δ*pchE* (“PA14 Δ*sid*”) and the mutant strain lacking both phenazines and siderophores (PA14 Δ*phz Δsid)* - as well as *B. glumae* wild-type and a mutant lacking the toxoflavin biosynthesis gene *toxA* (*B. glumae* Δ*toxA*). The effective Stern Volmer constants reported in the main-text represent the slope of linear least-squares fits to these curves. The bottom right panel gives the optical densities (500 nm) of the cultures from which the supernatants derive. Mutant strains did not differ drastically from their respective wild-types in end-point density. Markers indicate distinct replicates.

**Figure S3: O_2_NS Stern-Volmer plots for the strains presented in Figure 1D.** In addition to the strains presented in the main text, we tested supernatants from two PA14 mutants - a double siderophore knockout PA14 Δ*pvdA* Δ*pchE* (“PA14 Δ*sid*”) and the mutant strain lacking both phenazines and siderophores (PA14 Δ*phz* Δ*sid*) - as well as *B. glumae* wild-type and a mutant lacking the toxoflavin biosynthesis gene *toxA* (*B. glumae* Δ*toxA*). The effective Stern Volmer constants reported in the main-text represent the slope of linear least-squares fits to these curves. The bottom right panel gives the optical densities (500 nm) of the cultures from which the supernatants derive. Mutant strains did not differ drastically from their respective wild-types in end-point density. These experiments were conducted on a separate day and with separate cultures from those in Figure S3. Markers indicate distinct replicates.

**Figure S4: Fit values for effective Stern-Volmer constants with respect to RTDP and O_2_NS for strains presented in Figures S2 and S3.** Bar plots give the mean and 95% confidence interval on the mean slope of the Stern-Volmer plots presented in those figures. Note that the Stern-Volmer plots give only the slope of the curve and not its Y-intercept.

**Figure S5: RTDP Stern-Volmer plots for the chemicals presented in Figure 1I.** The Stern Volmer constants reported in the main-text represent the slope of linear least-squares fits to these curves. Markers indicate distinct replicates.

**Figure S6: O_2_NS Stern-Volmer plots for the chemicals presented in Figure 1J.** The Stern Volmer constants reported in the main-text represent the slope of linear least-squares fits to these curves. Markers indicate distinct replicates. The highest concentration of PMS exhibited strongly super-linear quenching of O_2_NS. We therefore excluded the highest PMS concentration (above the dashed gray line) when fitting *K_sv_*.

**Figure S7: RTDP Stern-Volmer plots for the strains presented in Figure 2A.** The Stern-Volmer constants reported in the main-text represent the slope of linear least-squares fits to these curves. In contrast to Figure 1C-D, strains were grown in glucose minimal media supplemented with amino acids, which may explain the difference in quenching from *E. coli* supernatants.

**Figure S8: O_2_NS Stern-Volmer plots for the strains presented in Figure 2B.** The Stern-Volmer constants reported in the main-text represent the slope of linear least-squares fits to these curves. In contrast to Figure 1C-D, strains were grown in glucose minimal media supplemented with amino acids, which may explain the difference in quenching from *E. coli* supernatants.

